## Supplementary Figures and Tables for "Characterizing the variation in chromosome structure ensembles in the context of the nuclear microenvironment"

This supplementary file contains:

Supplementary Tables 1 and 2

Supplementary Figures S1 to S10

| | **IMR90** | **K562** | **A549** |
|---|---|---|---|
| **H3K4Me1** | ENCFF812CWF | ENCFF352HXD | ENCFF269UHZ |
| **H3K4Me2** | ENCFF842IIU | ENCFF446FUS | ENCFF143DSJ |
| **H3K4Me3** | ENCFF021CLP | ENCFF855ZMQ | ENCFF756VHC |
| **H3K9Ac** | ENCFF733ZHB | ENCFF149MXA | ENCFF497JCB |
| **H3K27Ac** | ENCFF064XTS | ENCFF121RHF | ENCFF921QFJ |
| **H4K20Me1** | ENCFF951EYC | ENCFF923YUN | ENCFF841QBW |
| **H3K36Me3** | ENCFF994SJM | ENCFF880HKV | ENCFF347QGE |
| **H3K27Me3** | ENCFF547BFL | ENCFF392ZKG | ENCFF979OSH |
| **H3K9Me3** | ENCFF084CZB | ENCFF155UQU | ENCFF089MBN |
| **H2A.Z** | ENCFF640IYV | ENCFF874SMO | ENCFF159DMB |
| **DNase** | ENCFF530KLY | ENCFF257HEE | ENCFF539TQP |
| **RAD21** | ENCFF895JAW | ENCFF057JFH | - |

Supplementary Table 1. ENCODE accession IDs for the eleven epigenetic marks from three different cell-types.

| | **Active State 1** | **Active State 2** | **Inactive Poised** | **Repressive Polycomb** | **Heterochromatin** |
|---|---|---|---|---|---|
| **Active State 1** | 0.8 \| 1.3 | 0.75 \| 1.3 | 0.85 \| 1.3 | 0.9 \| 1.3 | 0.95 \| 1.3 |
| **Active State 2** | 0.75 \| 1.3 | 0.7 \| 1.3 | 0.75 \| 1.3 | 0.8 \| 1.3 | 0.85 \| 1.3 |
| **Inactive Poised** | 0.85 \| 1.3 | 0.75 \| 1.3 | 1.0 \| 1.35 | 1.0 \| 1.35 | 1.0 \| 1.35 |
| **Repressive Polycomb** | 0.9 \| 1.3 | 0.8 \| 1.3 | 1.0 \| 1.35 | 1.3 \| 1.4 | 1.1 \| 1.35 |
| **Heterochromatin** | 0.95 \| 1.3 | 0.85 \| 1.3 | 1.0 \| 1.35 | 1.1 \| 1.35 | 1.4 \| 1.4 |

Supplementary Table 2. Parameters values for Lennard-Jones Potential $U_P(r)$ used in the polymer model. Each entry represents the interaction strength and distance cutoff (separated by |) between beads of that two specific types. Energy values and distance cutoff values are represented in terms of $k_B T$ and $\sigma$ respectively.

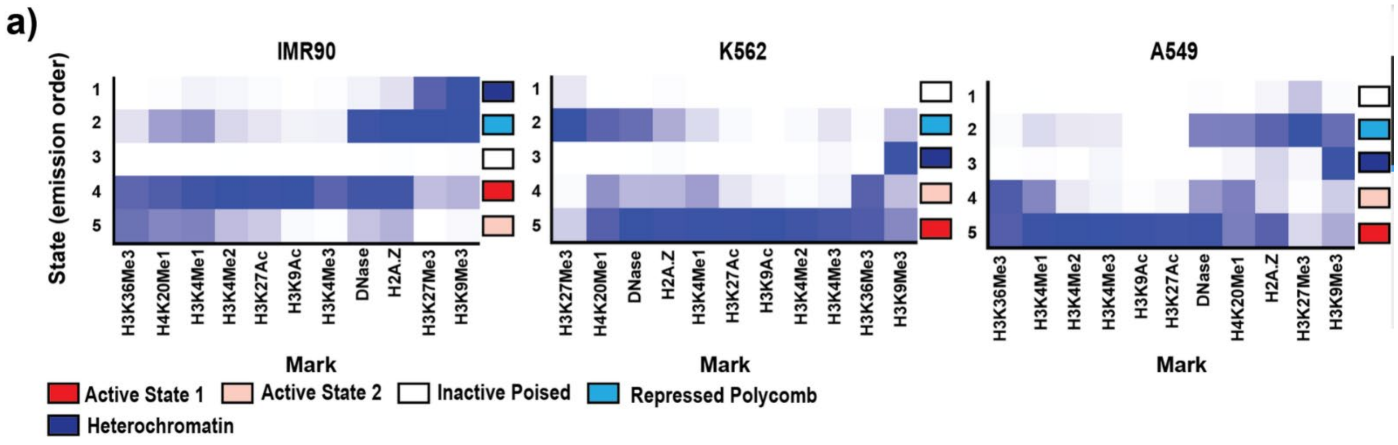


**Supplementary Figure 1**. Assignment of different chromatin states based on relative enrichment of different histone marks using ChromHMM tool for chr21:28-30 Mb genomic region from three different cell-types - IMR90, K562 and A549 at 30 Kb resolution.

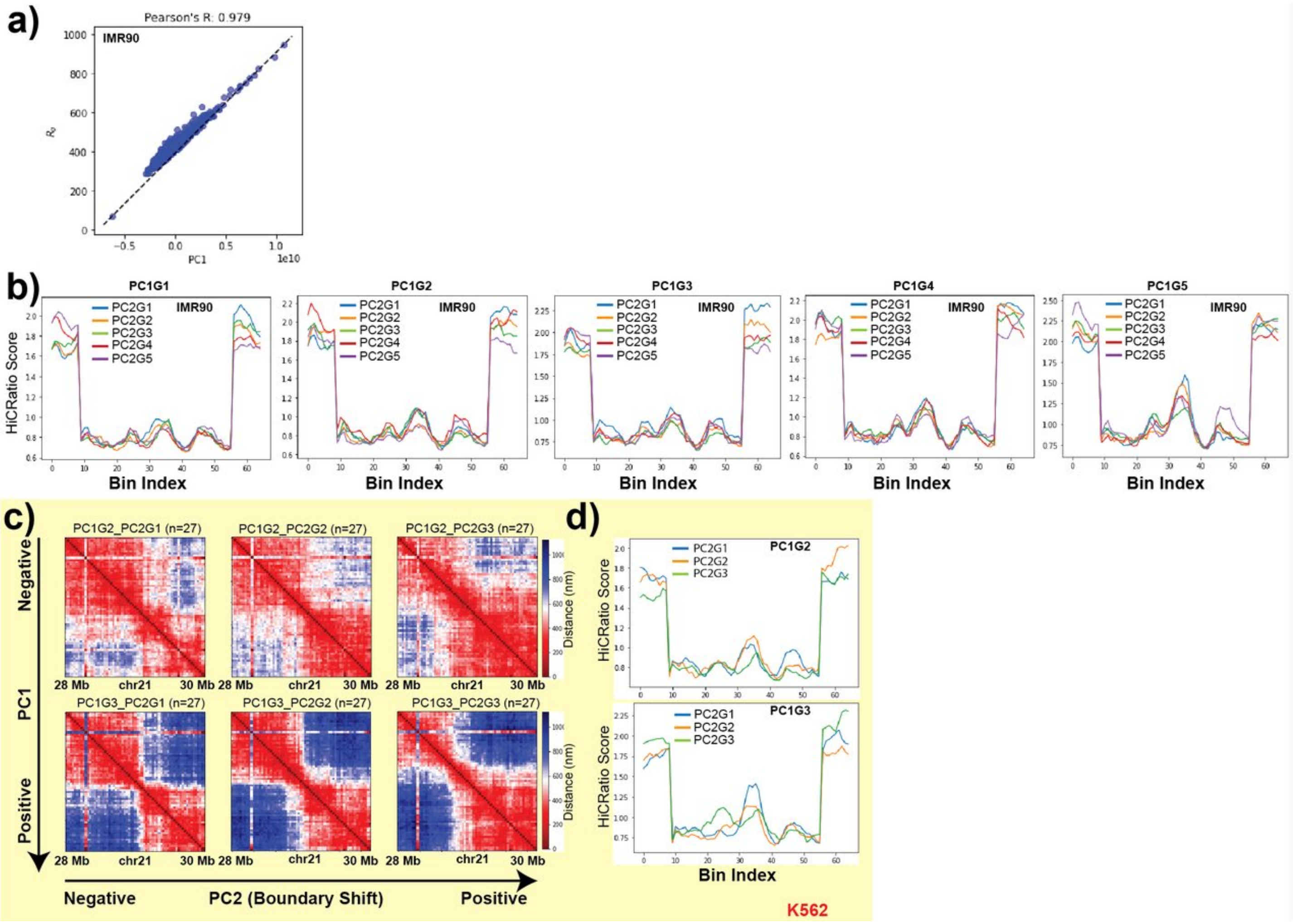


**Supplementary Figure 2**. a) Pearson's correlation between chromosome structural landscape PC1 values and radius of gyration ($R_g$) of single-cell structures (calculated from 3D position traces) for IMR90 chr21:28-30 Mb genomic region. b) HiCRatio score of PC2 ordered structures belonging to different PC1 groups for IMR90 chr21:28-30 Mb genomic region, calculated by HiCRatio approach (with a 300 kb window size). Here, PC1G1 refers to the group1 (G1; smallest $R_g$ structures) based on PC1. Within PC1G1, then PC2G1 means the group1 (G1) based on PC2. Successive higher numbered PC1 groups have higher $R_g$ structures. c) For K562 chr21:28-30 Mb genomic region, structures are first divided into three groups based on the PC1 ordering. Within each group, structures are further divided into three subgroups based on their PC2 ordering. To display the domain reorganization along PC2 in K562, averaged distance maps from only groups 2 and 3 from PC1 are shown here. d) For the same groups as in c, HiCRatio (with a 300 kb window size) scores of structures are shown. Colored lines represent HiCRatio scores from PC2 subgroups within two different PC1 subgroups (top = smaller $R_g$ and bottom = larger $R_g$)

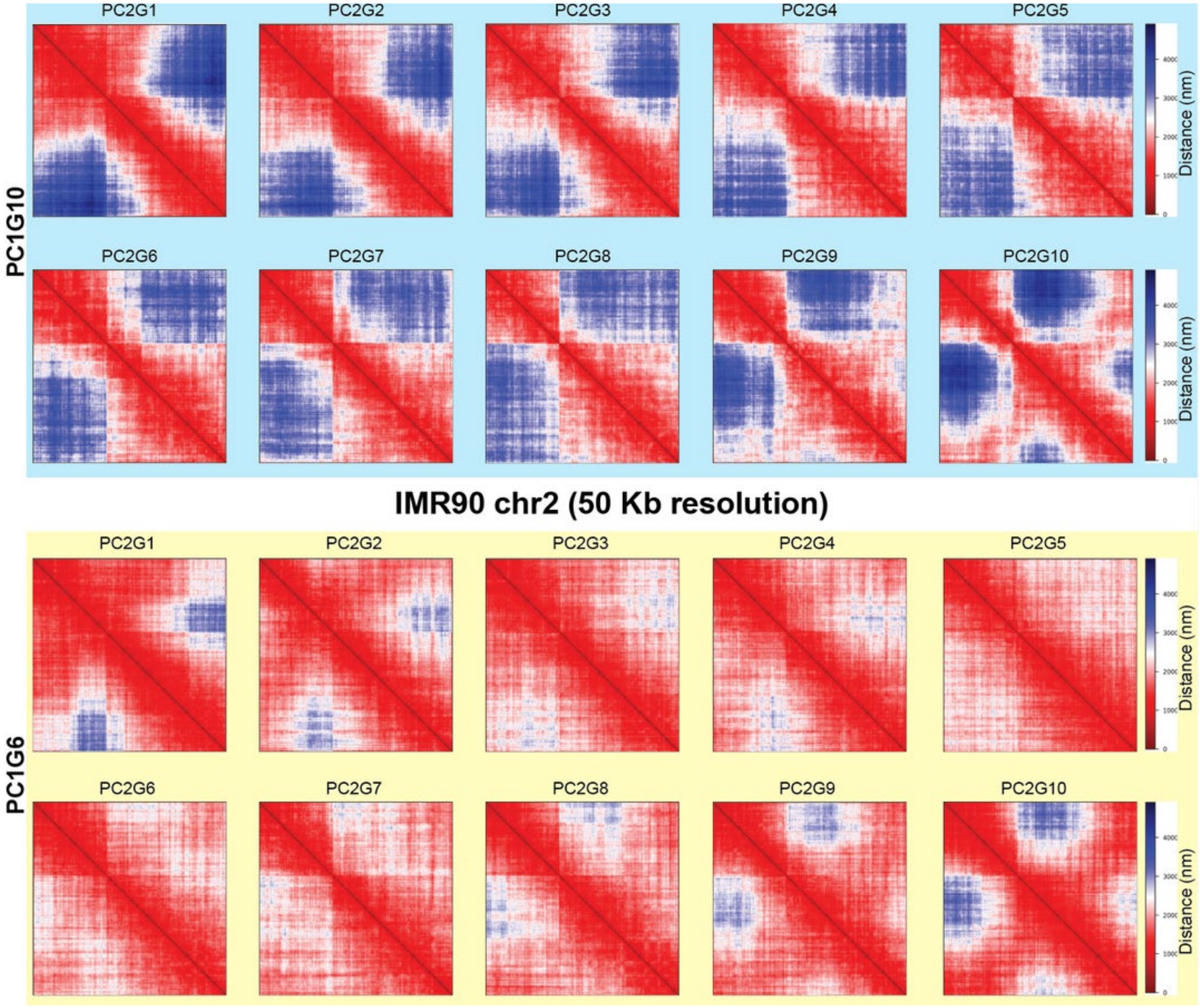


**Supplementary Figure 3**. For IMR90 chr2 structure data, structures are first divided into ten groups based on the PC1 ordering. Within each group, structures are further divided into ten subgroups based on their PC2 ordering. To display the domain reorganization phenomenon along PC2 in IMR90 chr2, only PC1 subgroups 6 (lower $R_g$) and 10 (higher $R_g$) are shown here. Heatmaps display the average distances for structures within each PC2 subgroup in each PC1 subgroup.

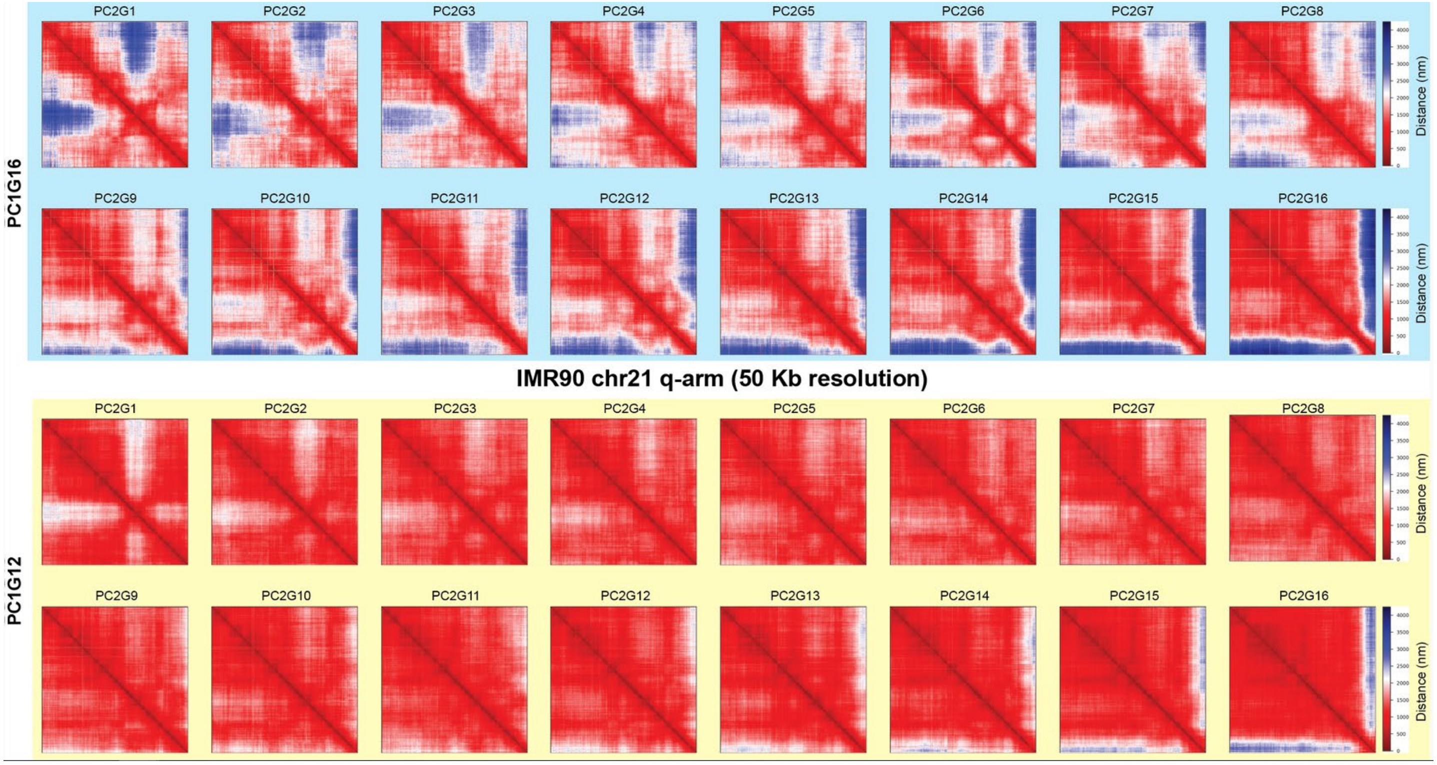


**Supplementary Figure 4**. For IMR90 chr21 entire q-arm structure data, structures are first divided into sixteen groups based on the PC1 ordering. Within each group, structures are further divided into sixteen subgroups based on their PC2 ordering. To display the domain reorganization phenomenon along PC2 in IMR90 chr21, only PC1 subgroups 12 (lower $R_g$) and 16 (higher $R_g$) are shown here. Heatmaps display the average distances for structures within each PC2 subgroup in each PC1 subgroup.

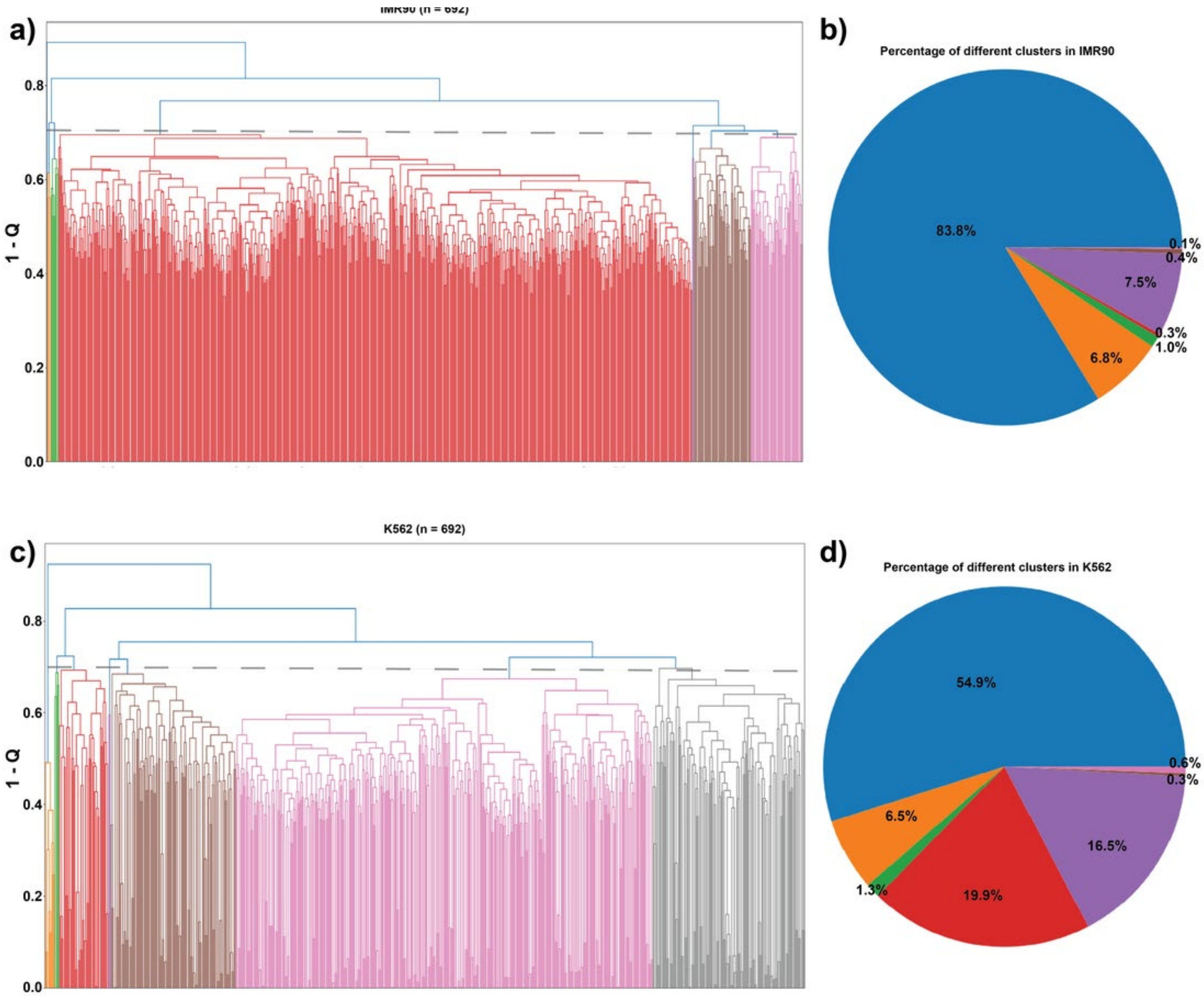


**Supplementary Figure 5**. a,c) Hierarchical clustering of IMR90 (a) and K562 (c) structures for chr21:28-29.5 Mb genomic segment, based on $Q$ similarity metric as described in Cheng *et al*. (1). To assign structures to the different clusters, the trees are cut at 0.7. b,d) Proportion of structures in different clusters obtained from IMR90 (b) and K562 (d) structural ensembles.

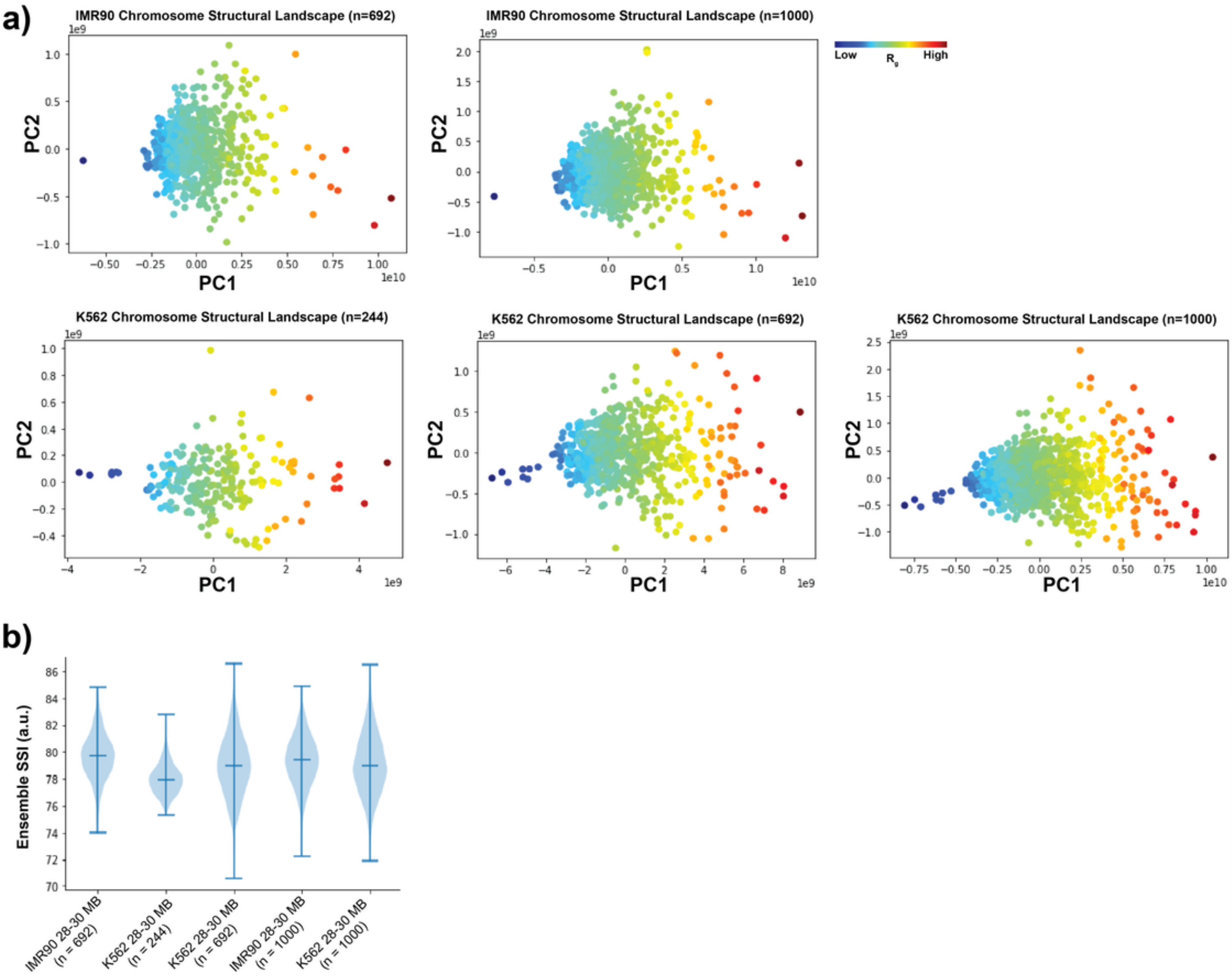


**Supplementary Figure 6**. a) Chromosome structural landscape of IMR90 and K562 chr21:28-30 Mb chromatin regions constructed for different numbers of cells. Points are colored according to the radius of gyration ($R_g$) of the corresponding structures (dark blue – smaller $R_g$ and dark red – larger $R_g$). b) Chromosome structural ensemble SSI distributions for chr21:28-30 Mb segment from IMR90 and K562 for different number of cells (shown as n). Here, for resampling 200 single-cell structures are selected randomly each time.

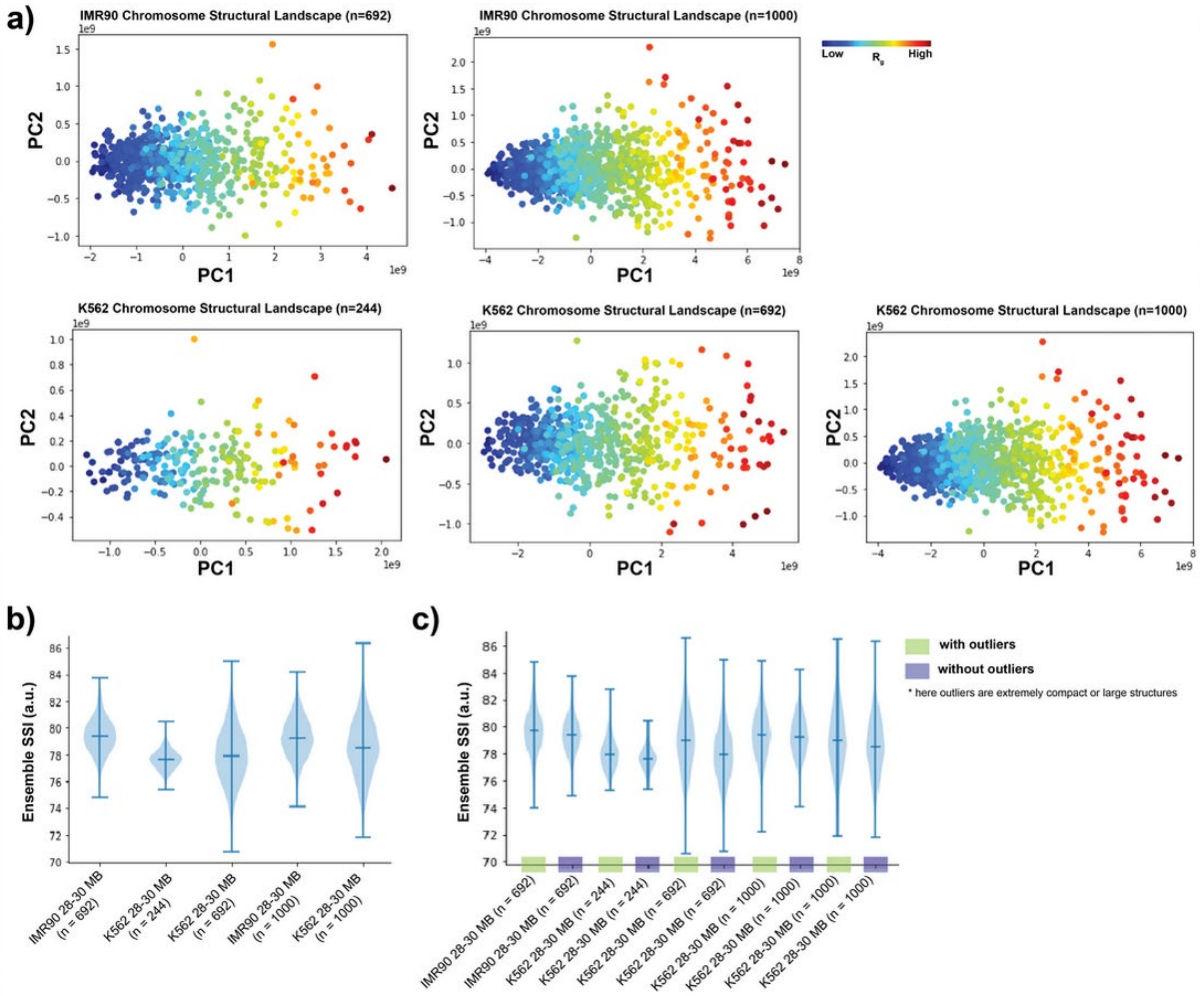


**Supplementary Figure 7**. a) Chromosome structural landscape of IMR90 and K562 chr21:28-30 Mb chromatin regions constructed for different numbers of cells after removing extremely compact or large conformations. Points are colored according to the radius of gyration ($R_g$) of the corresponding structures (dark blue – smaller $R_g$ and dark red – larger $R_g$). b,c) Chromosome structural ensemble SSI distributions for chr21:28-30 Mb segment from IMR90 and K562 for different number of cells, before (c) or after (b) removing extremely compact or large conformations. Here, for resampling 200 single-cell structures are selected randomly each time.

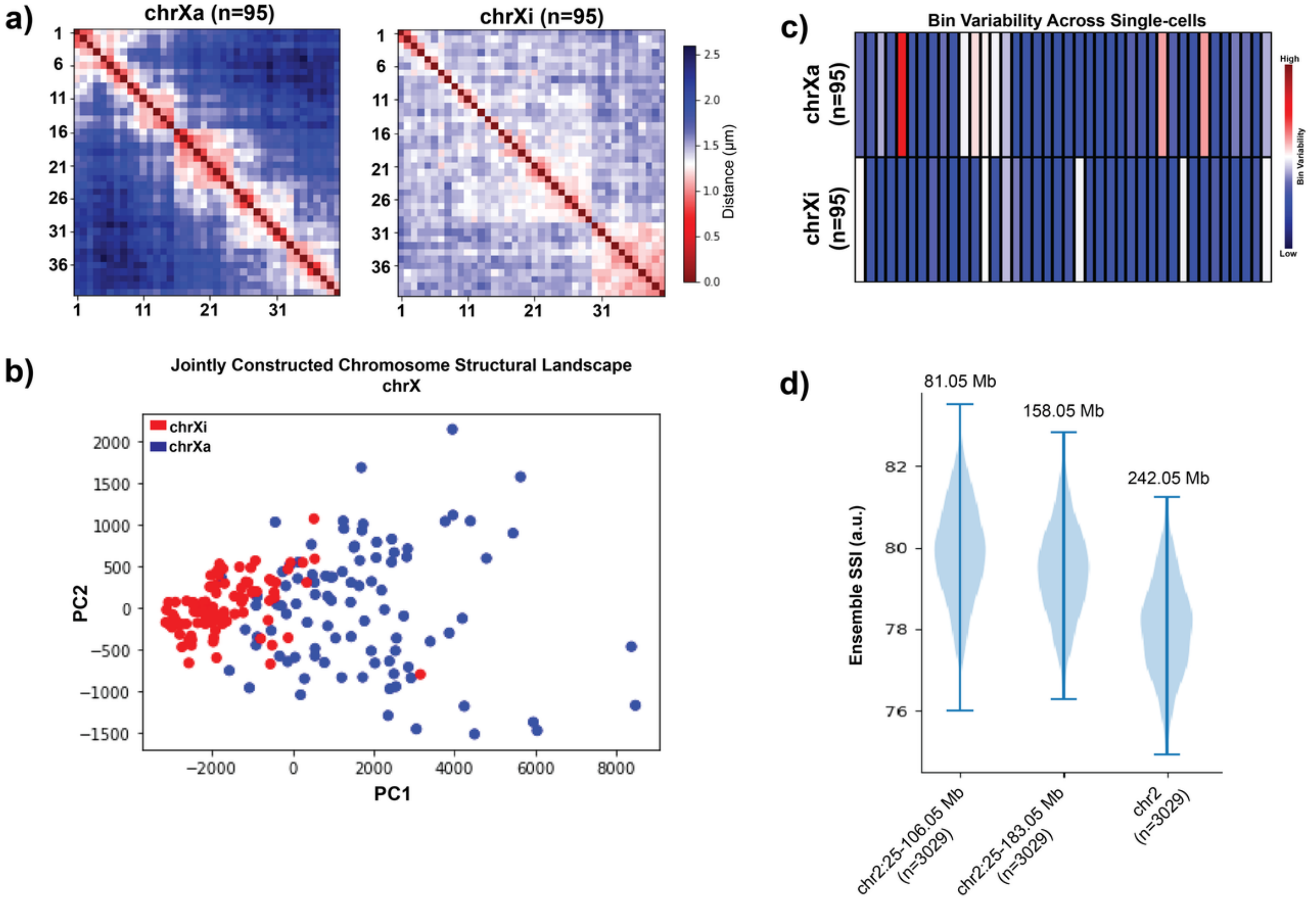


**Supplementary Figure 8**. a) Population-averaged distance maps of active and inactive chrX from IMR90 cell at TAD resolution. b) Jointly constructed chromosome structural landscape of active and inactive chrX from IMR90 cell. c) Bin Variability (BSI standard deviation) of active and inactive chrX, where a higher value represents a higher level of structure variation for that bin across the population and vice versa. d) Chromosome structural ensemble SSI distributions for chr2 segments of varying length. Here, for resampling 300 single-cell structures are selected randomly each time.

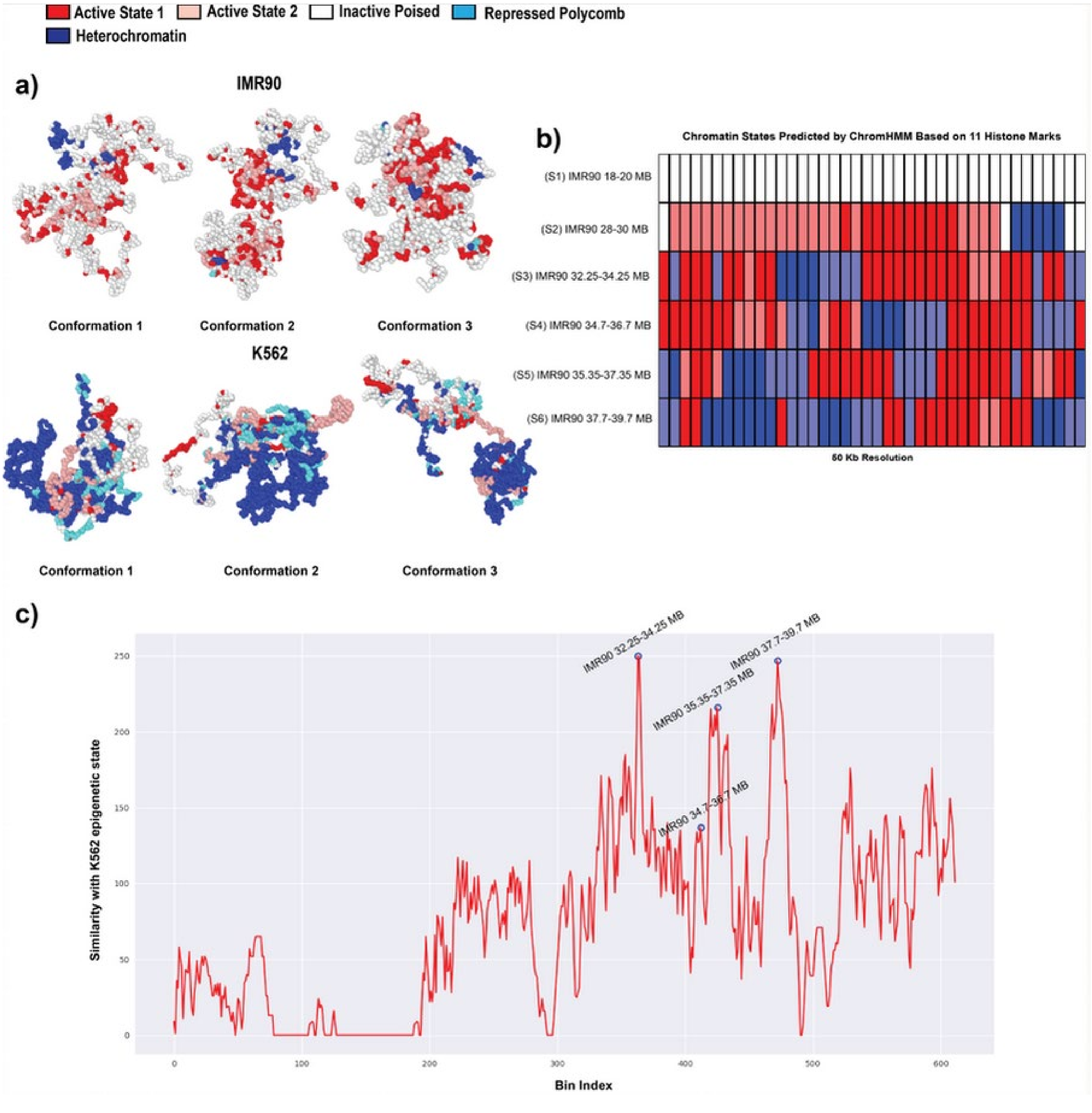


**Supplementary Figure 9**. a) Representative polymer models of IMR90 and K562 for chr21:28-29.5 Mb genomic segment, obtained from different time points of the simulation. The color of the beads shows corresponding epigenetic states. b) Chromatin states of six different 2 Mb spanning genomic regions of IMR90 chr21 at 50 kb resolution. The chromatin states are inferred based on 11 different histone marks for each cell-type by ChromHMM tool. c) Epigenetic state similarity of different 2 Mb spanning genomic regions of IMR90 chr21 q-arm with the K562 28-30 Mb region using binary comparison. Here, a higher comparison value shows higher similarity with the K562 epigenetic state.

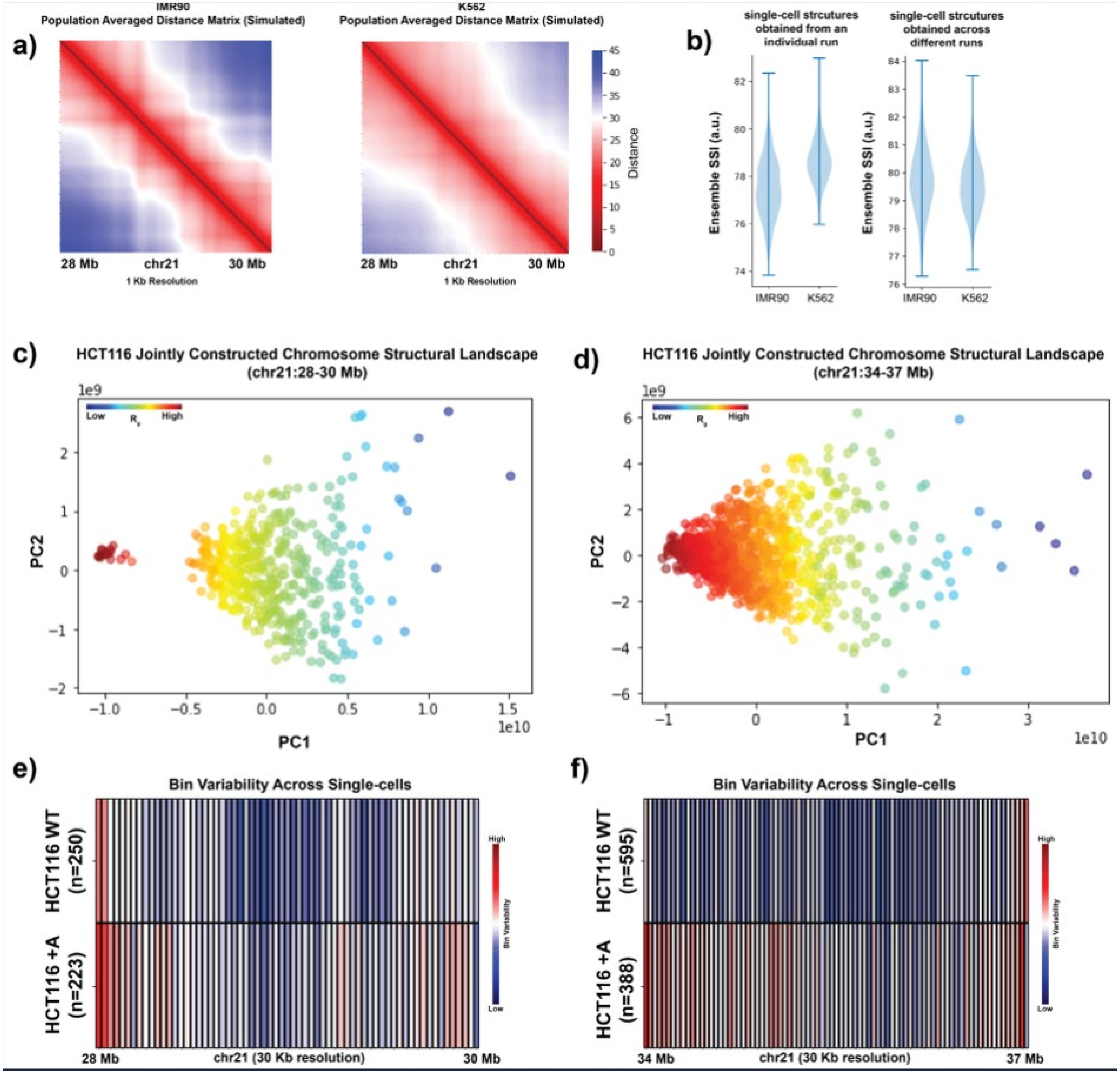


**Supplementary Figure 10**. a) Population-averaged distance maps of IMR90 and K562 chr21:28-30 Mb genomic region from loop extrusion polymer simulations (see Methods) at 1 kb resolution. b) Chromosome structural ensemble SSI distributions for chr21:28-30 Mb from loop extrusion simulated IMR90 and K562. For resampling, 500 single-cell structures are selected randomly each time from a single run (left) or across multiple runs (right). c,d) Jointly constructed chromosome structural landscapes of chr21:28-30 Mb (c) or chr21:34-37 Mb (d) of HCT116 for WT and +Auxin conditions. Points are colored according to the radius of gyration ($R_g$) of the corresponding structures (dark blue – smaller $R_g$ and dark red – larger $R_g$). e) Bin Variability of chr21:28-30 Mb genomic region for wild-type (WT) and cohesin-depleted (+Auxin) conditions. f) Bin Variability of chr21:34-37 Mb genomic region for wild-type (WT) and cohesin-depleted (+Auxin) conditions.
